# From genotype to antibiotic susceptibility phenotype in Enterobacteriaceae: a clinical perspective

**DOI:** 10.1101/503607

**Authors:** Etienne Ruppé, Abdessalam Cherkaoui, Myriam Girard, Stéphane Schicklin, Vladimir Lazarevic, Jacques Schrenzel

**Author notes:** Current affiliation: INSERM, IAME, UMR 1137; Université Paris Diderot, IAME, UMR 1137, Sorbonne Paris Cité and AP-HP, Hôpital Bichat, Laboratoire de Bactériologie, F-75018 Paris, France. Corresponding author: Etienne RUPPE (PharmD, PhD), Laboratoire de Bactériologie, Hôpital Bichat-Claude Bernard, 46 rue Henri Huchard, 75018 Paris, France.

## Abstract

Predicting the antibiotic susceptibility phenotype from genomic data is challenging, especially for some specific antibiotics in Enterobacteriaceae. Here we aimed to assess the performance of whole genomic sequencing (WGS) for predicting the antibiotic susceptibility in various Enterobacteriaceae species using the detection of antibiotic resistance genes (ARGs), specific mutations and a knowledge-based decision algorithm. We sequenced (Illumina MiSeq 2×250b) 187 clinical isolates from species possessing (n=98) or not (n=89) an intrinsic AmpC-type cephalosporinase. Antibiotic susceptibility was performed by the disc diffusion method. Reads were assembled by A5-miseq and ARGs were identified from the ResFinder database using Diamond. Mutations on GyrA and ParC topoisomerases were studied. We assessed the prediction rates for amoxicillin, co-amoxiclav, piperacillin, piperacillin-tazobactam, ceftazidime, cefepime, meropenem, amikacin, gentamicin and fluoroquinolones. A total of 1,870 isolate/antibiotic combinations were considered. In 295 cases (15.8%) no attempt of prediction was made. Among the 1,575 attempts, 1,537 (97.6%) were correct (1,011 for predicting susceptibility and 526 for predicting resistance), 15 (0.9%) were major errors (MEs) and 23 (1.5%) were very major errors (VMEs). The concordance rates were similar between non-AmpC and AmpC-producing Enterobacteriaceae (907/935 [97.0%] vs 630/640 [98.4%], Chi2 test p=0.07), but more VMEs were observed in non-AmpC producing strains than in those producing an AmpC (20/935 [2.1%] vs 3/640 [0.5%], Chi2 test p=0.007). The majority of VMEs were putatively due to the overexpression of chromosomal genes. In conclusion, the inference of antibiotic susceptibility from genomic data showed good performances for non-AmpC and AmpC-producing Enterobacteriaceae species.

## Introduction

Whole genome sequencing (WGS) of bacterial strains has now become the gold standard for the identification of antibiotic resistance determinants that includes antibiotic resistance genes (ARGs) and intrinsic genes in which mutational events can lead to antibiotic resistance. Yet, the *in silico* translation from genotype to phenotype may be challenging because it relies on the quality and exhaustiveness of the available knowledge about the genomic determinants of resistance. First, the ARG database must be exhaustive so that no ARG shall be missed. Several ARG databases have been released over the last decade (1), the most popular ones being ResFinder (2) and CARD (3). To date, there is no consensus on which database should be used for inferring antibiotic susceptibility phenotype from WGS data. Then, the resistance pattern conferred by the ARGs needs to be known, which is not the case for some variants that have not been experimentally tested (*i.e.* mutations of unknown phenotypic significance). Of note, no database includes phenotypic data associated with the ARG sequence and the resistance phenotype conferred by the presence of an ARG must be inferred from literature. Even more incomplete are the data relative to the mutational events associated with antibiotic resistance such as those leading to a decreased affinity for the antibiotic (e.g. mutations in the topoisomerase for fluoroquinolone resistance), an increased expression of an intrinsic resistance gene (e.g. *bla*_AmpC_ in Enterobacteriaceae) and/or a decreased expression of a gene (e.g. *oprD* in *Pseudomonas aeruginosa*) alone or in combination. Unlike acquired ARGs that have been thoroughly collected, data linking specific mutational events with a resistance phenotype are lacking, thereby introducing some caveats in the genotype-to-phenotype prediction for some bacteria-antibiotics combinations.

The link between the content of antibiotic resistance determinants (referred to as the “genotype”) and the antibiotic resistance profile (the “phenotype”) has been assessed for *Staphylococcus aureus* (4–7), *Escherichia coli* (8–11), *Shigella sonnei* (12), *Klebsiella pneumoniae* (9, 13), *P. aeruginosa* (14, 15) and *Mycobacterium tuberculosis* (16). As for *E. coli/S. sonnei* and *K. pneumoniae*, the performances of WGS to predict the susceptibility was excellent for beta-lactams, fluoroquinolones and aminoglycosides (17). In those species, resistance to antibiotics is driven by the acquisition of ARGs, which can be easily addressed by WGS. Nonetheless, in some situations, mutational events need to be analyzed, such as mutations in the topoisomerases for the resistance to fluoroquinolones (18), or mutations in the promoter region of the cephalosporinase-encoding gene *ampC* in *E. coli* (19). Some other Enterobacteriaceae species harbor an inducible, AmpC-type cephalosporinase that, when expressed at a basal level, confers a low-level resistance to beta-lactams. The *ampC*-repression mechanism involves AmpD (an amidase) and AmpR (a transcriptional regulator encoded by the gene located right upstream of *ampC*). A few studies have reported that the induction of *ampC* expression could be due to mutational events compromising the functions of AmpD and/or AmpR (20). The resulting AmpC overproduction causes resistance to all beta-lactams except the so-called fourth generation cephalosporins (cefepime) and carbapenems. Still, the census of genetic events leading to an overproduction of AmpC in the context of inferring antibiotic susceptibility from WGS data has not been performed.

The use of WGS to infer antibiotic susceptibility could be translated in the clinical setting through clinical metagenomics (CMg), which refers to the metagenomic sequencing the nucleic acid from clinical samples in order to obtain information of clinical relevance. CMg is an emerging field that could transform the way infectious diseases are currently diagnosed. Indeed, CMg applied to various clinical samples has shown promising results in the identification of bacterial pathogens, including the hardly culturable ones. The identification of ARGs and the inference of antibiotic susceptibility should be the next step to achieve for CMg in order to provide a complete bacteriological analysis. CMg can readily be performed in 48-72h using Illumina sequencing platforms, which is a reasonable turn-around time when it comes to the bacteriological diagnostic of non-severe infections. As for severe infections such as hospital-acquired pneumonia (HAP), an early identification of infection-causing bacteria and the inference of its antibiotic susceptibility pattern should improve the patient’s care in providing an optimized antibiotic regimen. In this perspective, the fast turn-around time offered by Oxford Nanopore Technologies sequencers makes the use of CMg in severe infections such as HAP possible (21).

The main bacteria causing HAP are *S. aureus, P. aeruginosa* and Enterobacteriaceae (22). As the antibiotic susceptibility inference by WGS has already been studied for *P. aeruginosa* and *S. aureus* but only for two Enterobacteriaceae species (*E. coli* and *K. pneumoniae*), here we aimed at assessing the performances of WGS to infer the susceptibility of various species of Enterobacteriaceae to the antibiotics commonly used in probabilistic therapy of HAP.

## Material and Methods

### Selection of the strains

A total of 187 Enterobacteriaceae strains from the bacteriology laboratory of the Geneva University Hospitals (HUG) have been analyzed for this project (Supplementary Figure 1). The strains have been selected according to the following criteria: (1) Enterobacteriaceae species underrepresented in genotype-to-phenotype studies (*i.e*. Enterobacteriaceae other than *E. coli* and *K. pneumoniae*), and (2) strains with an antibiotic susceptibility profile of interest (with the aim of recovering a high diversity of antibiotic susceptibility profiles). Only one strain per patient was selected. Strain identification was performed using matrix-assisted desorption ionization-time of flight mass spectrometry (MALDI-TOF MS; Maldi Biotyper compass, Bruker Daltonics, Bremen, Germany) according to the manufacturer’s instructions.

The antibiotic susceptibility testing was performed using the disk-diffusion test methods according to the European Committee on Antimicrobial Susceptibility Testing (EUCAST v6.0) methods. The antibiotics considered for this study were penicillins (amoxicillin, piperacillin), penicillins+beta-lactamases inhibitors (co-amoxiclav, piperacillin-tazobactam), cephalosporins (ceftazidime and cefepime), carbapenems (meropenem), aminoglycosides (gentamicin and amikacin) and fluoroquinolones (norfloxacin).

### DNA extraction and genome sequencing

Genomic DNA of each isolate was extracted from colonies grown overnight at 37°C on blood agar plates using the MagCore Genomic DNA Tissue Kit (RBC Bioscience, New Taipei City, Taiwan), as described previously (23). Purified DNA was sent to Fasteris (Plan-les-Ouates, Switzerland) for sequencing. The library was prepared using the Nextera XT DNA Sample Preparation Kit according to the Illumina (San Diego, CA) instructions, and was sequenced on an Illumina MiSeq with 2 × 250 cycles. Six sequencing runs were performed, each including 21 to 50 multiplexed samples.

### Bioinformatic analyses

The mean number of high quality (paired-end) reads per sequenced strain was 603,139 and the mean coverage was 27x. The reads were processed following the pipeline depicted in the Supplementary Figure 2. Briefly, raw reads were trimmed and quality-filtered using Trimmomatic (Q20 over a 4-nucleotide window) (24). The quality of the reads was assessed by FastQC before and after the quality filter. Species identity was verified using MetaPhlAn2 (25). Quality reads were assembled using the A5-miseq assembler (26) (version 20160825:243). Genes were predicted and annotated using PROKKA (27). Antibiotic resistance genes were sought using Diamond (28) and the ResFinder database, accessed October 2017 (2). The threshold for ARG positive detection was arbitrarily set at an h-value (defined as the product of the amino acid identity and the query coverage) ≥0.64. The amino-acid sequences of TEM and SHV beta-lactamases were respectively aligned with those of TEM-1 and SHV-1 in order to identify putative mutations and to assign the variant number according to the numbering scheme of the Lahey Clinic (now hosted by the NCBI). The amino acid sequences of the chromosomal genes associated with quinolone resistance (*gyrA* and *parC*) were aligned to the homologous protein sequence from the corresponding species (obtained from the NCBI reference genome) using MUSCLE (29). The *ampC* promoter region of *E. coli* was also analyzed for the presence of mutations (19). The results of the assembly process are shown in the Supplementary Figure 3. The genome sizes, estimated as the sum of contigs, varied from 3,729,115bp (*Morganella morganii* 123) to 6,417,271bp (*Klebsiella oxytoca* 94). The distribution of the genome sizes according to the species is depicted in the Supplementary Figure 4. Figures were generated using R v3.4.2 and the ggplot2 package. Colors were picked from colorBrewer.org.

### Genotype to phenotype inference

To determine links between genotype and phenotype, we applied specific rules related to the species, the presence of ARGs and the antibiotic (Table 1). We assumed that all the ARGs found in the strains were expressed except for some chromosomal ARGs (Table 1). In some cases, we considered that no inference could be performed because of the lack of data regarding the expression level of the ARG (e.g. *bla*_TEM-1_ and co-amoxiclav/piperacillin-tazobactam), and therefore no antibiotic-resistance prediction was attempted. A very major error (VME) was defined as inferring susceptibility from genomic data while the strain was actually resistant by phenotypic tests. A major error (ME) was defined as inferring resistance from genomic data while the strain was actually susceptible by phenotypic tests. In case of VMEs for beta-lactams inference, other beta-lactamases were searched in the ResFinderFG database (https://cge.cbs.dtu.dk/services/ResFinderFG/). The spectrum of activity of aminoglycosides-modifying enzymes (AMEs) was determined from Ramirez *et al* (30).

**Table 1:** Interpretation rules applied in this work for inferring an antibiotic susceptibility pattern from genomic data.

|  |
| --- |
| **Penicillins and combination penicillins+beta-lactamases inhibitors**<br>- Species other than *Escherichia coli*: resistance to AMX when any beta-lactamase was identified.<br>- *E. coli*: AMX, AMC, PIP and TZP resistance when the combination -42C>T and -18G>A, and/or the -32T>A mutation was found in the *ampC* promoter region.<br>- Presence of narrow-spectrum TEM (e.g. TEM-1): no prediction of AMC susceptibility.<br>- AmpC-producing Enterobacteriaceae: no prediction for PIP or TZP when no beta-lactamase other than the intrinsic AmpC was identified.<br>- Presence of an acquired AmpC: resistance to AMX, AMC, PIP and TZP.<br>- Sole production of an extended-spectrum beta-lactamase (ESBL): resistance to AMX and PIP. No prediction of AMC and TZP.<br>- Intrinsic AmpC-producing species were considered to be resistant to AMX and AMC.<br>- Intrinsic penicillinase-producing species (*Klebsiella pneumoniae*, *Klebsiella oxytoca*, *Citrobacter koseri*) were |
| **Ceftazidime**<br>(other than OXA-48-like) - encoding gene was found.<br>For *E. coli*, strains were considered to be susceptible to CAZ when no -42C>T and -18G>A, and/or the -32T>A were found.<br>For other AmpC-species: strains were considered to be susceptible to CAZ when both AmpD and AmpR had no mutation compared to the reference AmpD and AmpR proteins, and no ESBL or carbapenemase (other |
| **Cefepime**<br>encoding gene was found. |
| **Meropenem**<br>Strains were considered to be susceptible to MEM when no carbapenemase-encoding gene was found. |
| **Gentamicin**<br>Strains were considered to be resistant to GEN when an AAC(3) or a 16S rRNA methylase-encoding gene was found. |
| **Amikacin**<br>Strains were considered to be resistant to AMK when an AAC(6')-I and/or an APH(III)-VI and/or a 16S rRNA methylase was found. |
| **Norfloxacin**<br>- at least one mutation was found in the quinolone resistant determining region (QRDR) of GyrA and/or ParC<br>- and/or an acquired Qnr was found.<br>Species with an intrinsic *qnr* gene (*Citrobacter freundii*, *Serratia marcescens*) were considered to be susceptible to NOR.<br>Species with an intrinsic OqxAB efflux system (*Klebsiella pneumoniae*, *Klebsiella oxytoca*, *Enterobacter aerogenes*, *Enterobacter cloacae*, *Serratia marcescens*) were considered to be susceptible to NOR. |
| **AMX:** amoxicillin; **AMC:** co-amoxiclav; **PIP:** piperacillin; **TZP:** piperacillin-tazobactam; **CAZ:** ceftazidime; **FEP:** cefepime; **MEM:** meropenem; **GEN:** gentamicin; **AMK:** amikacin; **NOR:** norfloxacin. |

## Results

### Genotype-to-phenotype inference

The analysis of the genomic sequence of 187 strains revealed 1,055 ARGs with 133 being unique. The distribution of the ARGs according to the antibiotic family they confer resistance to is depicted in the Figure 1. The main ARG families were beta-lactamases (n=225), Tet (efflux and protection, n=220) and AMEs (n=177). Overall, 1,870 isolate-antibiotic combinations were considered (Table 2). In 295 cases (15.8%) no attempt of antibiotic resistance prediction was made. Attempts were less frequent in AmpC-producing than in non-producing isolates (640/890 [71.9%] vs. 935/980 [95.4%, respectively, Chi2 test p=4.6e-44).

**Figure 1:**
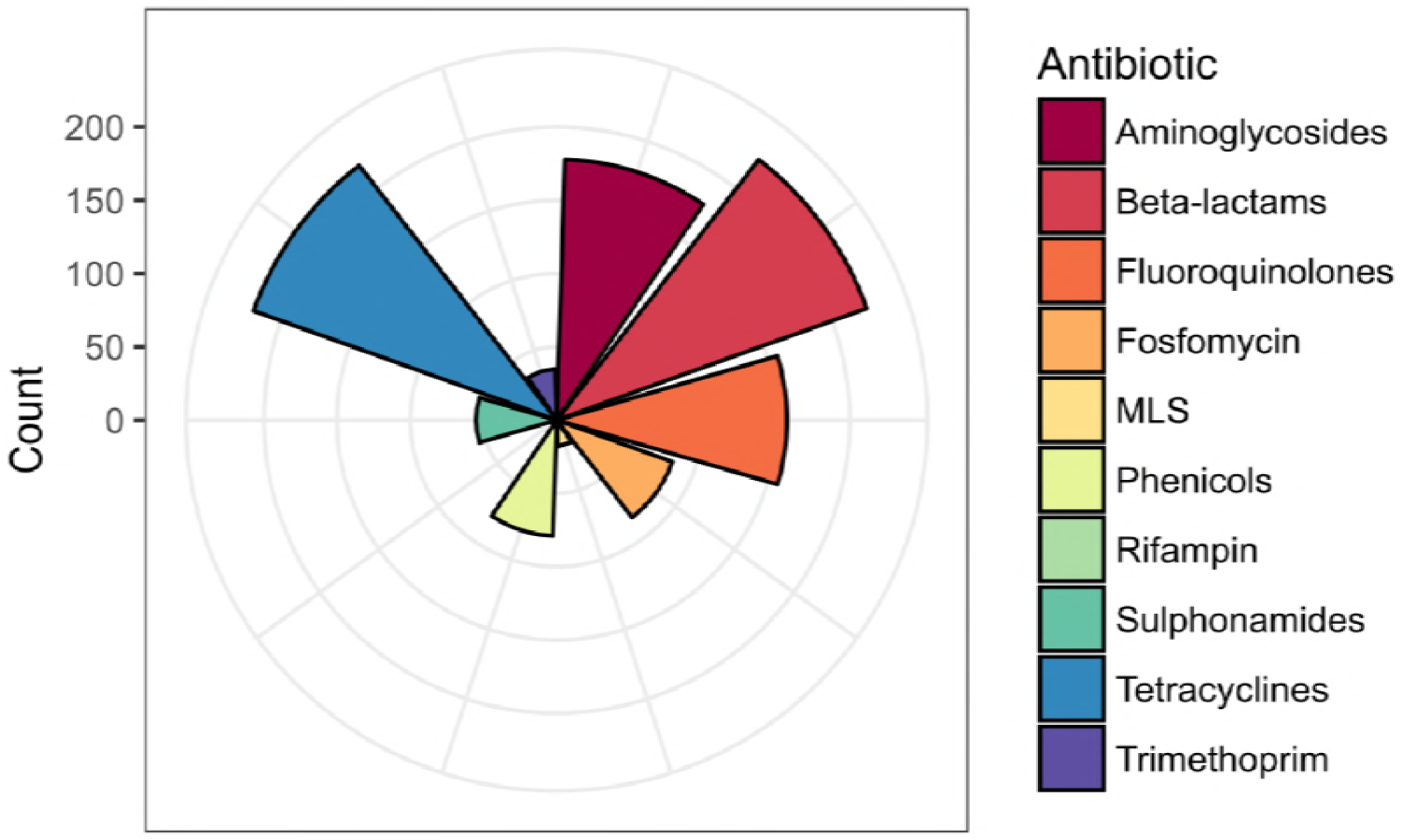
Distribution of the antibiotic resistance genes (ARGs) according to the antibiotic family they confer resistance to. MLS: macrolides, lincosamides and streptogramins.

**Table 2:** Results of the predictions.

| Prediction | Non-AmpC strains<br>(n=980 combinations) | AmpC-strains<br>(n=890 combinations) | All<br>(n=1,870 combinations) | Difference non-AmpC/AmpC |
| --- | --- | --- | --- | --- |
| Attempted | 935 (95.4%<sup>a</sup>) | 640 (71.9%<sup>a</sup>) | 1,575 (84.2%<sup>a</sup>) | p=4.6e-44 |
| Correct | 907 (92.2%<sup>b</sup>) | 630 (98.4%<sup>b</sup>) | 1,537 (97.6%<sup>b</sup>) | p=0.07 |
| Major error | 8 (0.9%<sup>b</sup>) | 7 (1.1%<sup>b</sup>) | 15 (0.9%<sup>b</sup>) | p=0.8 |
| Very major error | 20 (2.1%<sup>b</sup>) | 3 (0.5%<sup>b</sup>) | 23 (1.5%<sup>b</sup>) | p=0.007 |

<sup>a</sup>All combinations were considered. <sup>b</sup>only attempts were considered.

Among 1,575 attempts, 1,537 (97.6%) were correct (1,011 correctly predicting susceptibility and 526 correctly predicting resistance), 15 were MEs (0.9%) and 23 (1.5%) were VMEs. The correct prediction rates were similar between AmpC-producing and non-producing Enterobacteriaceae (630/640 [98.4%] vs 907/935 [97.0%], Chi2 test p=0.07), but more VMEs were observed in AmpC non-producing isolates (20/935 [2.1%] vs. 3/640 [0.5%], Chi2 test p=0.007). Detailed results of predictions by species are available in Table 3.

**Table 3:**
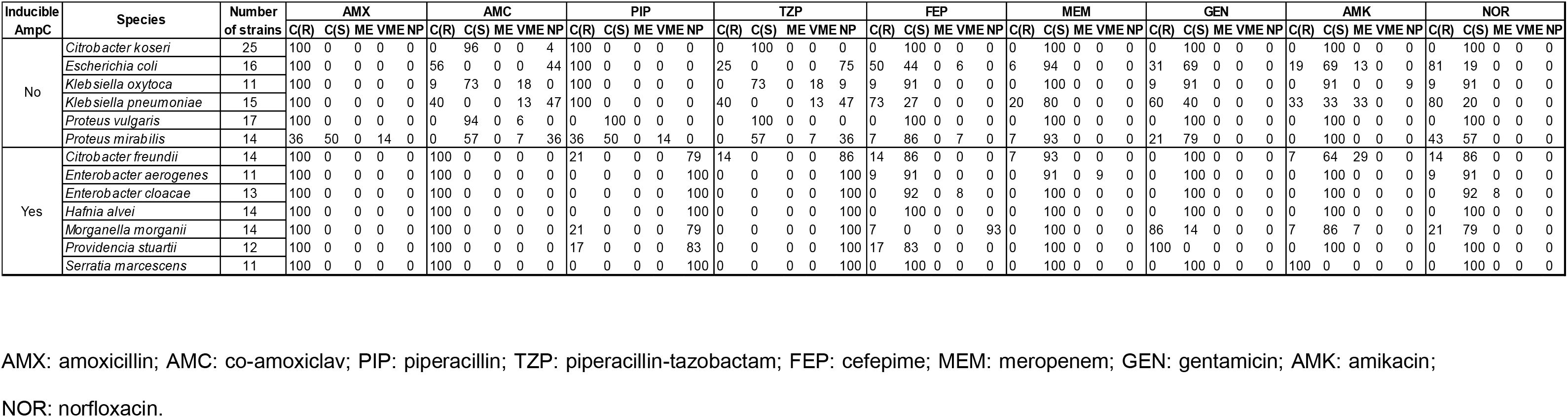
Detailed results of predictions (in percentages) by species.

### Penicillins and combination penicillins with beta-lactamase inhibitor

In total, 225 beta-lactamases were identified, 52 being unique (Supplementary Figure 5). The most common were CTX-M-15 and TEM-1. The correct rates of prediction were 98.9% for amoxicillin, 86.1% for co-amoxiclav, 55.6% for piperacillin and 37.4% for piperacillin-tazobactam (Figure 2). For penicillins, two VMEs (but no MEs) were found. The strain *P. mirabilis* 75 was resistant to penicillins, penicillins+beta-lactamases inhibitors and to extended-spectrum cephalosporins (but ceftriaxone) by phenotypic tests, however, no beta-lactamases or mutations on penicillin-binding proteins (PBP) were identified during the genome analysis (data not showed). The other VME, found in *P. mirabilis* 122, was due to the absence of a *bla*_HMS-1_ beta-lactamase from the current version of ResFinder (although it is listed in ResFinderFG). Finally, 6 VMEs for co-amoxiclav and 5 to piperacillin-tazobactam were likely due to the overexpression of chromosomal beta-lactamases (SHV-1, OXY-1, HUGA) and again to the peculiar phenotype of *P. mirabilis* 75.

**Figure 2:**
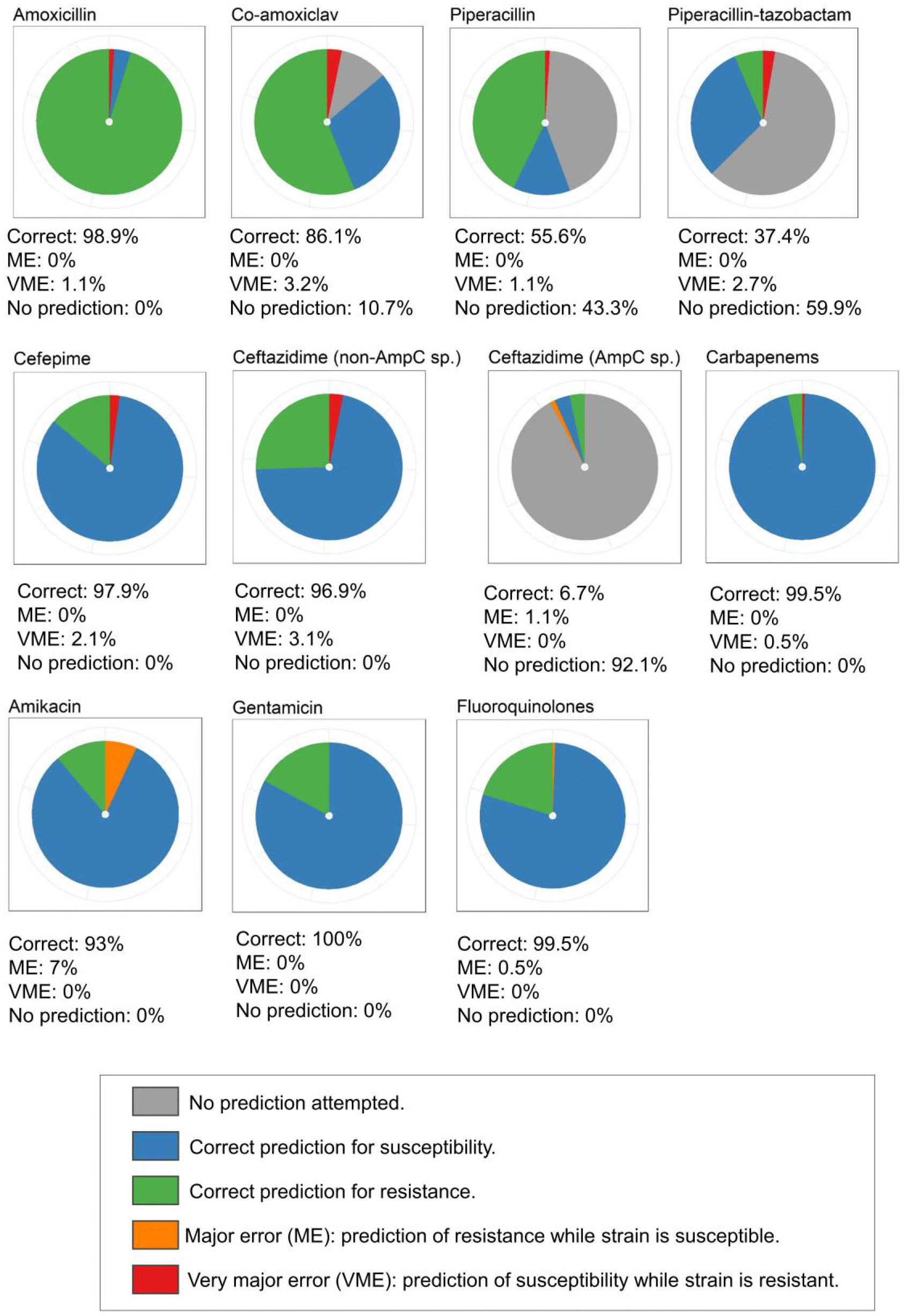
Performance of genotype to phenotype inference for the 187 strains of the study. AmpC sp: species harboring an intrinsic inducible AmpC-type cephalosporinase. Non-AmpC sp.: species not harboring an intrinsic inducible AmpC-type cephalosporinase.

### Extended spectrum cephalosporins (ceftazidime, cefepime) and carbapenems (meropenem)

Among the 225 beta-lactamases identified by WGS (Supplementary Figure 5), 28 were of extended-spectrum (26 CTX-M, 1 TEM-130 and 1 TEM-138). In all non-AmpC strains (n=98), the inference of ceftazidime susceptibility could be attempted. A correct inference was obtained in 95/98 (96.9%) cases (Figure 2), leaving 3 VMEs: two *K. pneumoniae* isolates with a putative overexpression of SHV-1 (31) and *P. mirabilis* 75. In AmpC strains, the inference of ceftazidime susceptibility could be attempted in only 7/89 cases (7.9%). The prediction was correct in 6/7 cases, with one major error (an *Enterobacter aerogenes* with a truncated *ampD* gene, yet susceptible to ceftazidime).

For cefepime and meropenem, the prediction could be performed for all strains. A correct inference was achieved in 97.9% (183/187) of strains (Figure 2). Four VMEs were observed for cefepime, including *P. mirabilis* 75 (see explanation above), *E. coli* 53 (combination of a CMY-42 and a OXA-181, both not being reported to alter cefepime susceptibility), *Enterobacter cloacae* 5 and *E. aerogenes* 240 (with likely combined overexpression of their chromosomal AmpC and decreased outer membrane permeability). As for meropenem, 5 carbapenemases were identified: 2 OXA-48, 1 OXA-181, 1 NDM-1 and 1 NDM-5. Only one VME was found for the same *E. aerogenes* 240 and a correct inference could be performed in 99.5% (186/187) strains (Figure 2).

In all, we identified 10 strains with at least a VME (with strains *P. mirabilis* 75 having 6 VMEs) and 1 with one ME, leaving 177/187 strains (94.7%) with a correct prediction for the tested beta-lactams.

### Aminoglycosides

We studied genotype-phenotype associations for two aminoglycoside antibiotics, gentamicin and amikacin. Resistance to aminoglycosides is due to the acquisition of AMEs (phosphotransferases [APH], nucleotidyltransferases [ANT] and acetyltransferases [AAC]) and 16S rRNA methyltransferases (e.g. Arm, RmtB). We identified 177 AMEs and one 16S rRNA methyltransferase, with 24 being unique (Supplementary Figure 6). For gentamicin, all predictions were correct (Figure 2). For amikacin, only MEs were observed (Figure 2). Indeed, 13 strains (5 *K. pneumoniae*, 4 *C. freundii*, 2 *E. coli*, 1 *K. oxytoca* and 1 *M. morganii*) bearing AAC(6’)-Ib-cr and/or an AAC(6’)-If were reported to be susceptible while AAC(6’)-I enzymes confer resistance to amikacin (32). No VMEs were observed for amikacin.

### Fluoroquinolones

Fluoroquinolone resistance occurs through mutations in a specific region of the topoisomerases (the quinolone resistance determining region, QRDR) and/or the acquisition of plasmid-mediated quinolone resistance (PMQR) elements such as Qnr. The distribution of GyrA and ParC topoisomerases mutations in the QRDR is depicted in the Supplementary Figure 7. A total of 38 strains had non-synonymous mutations in in the QRDR of *gyrA* and/or *parC* genes. Any amino acid substitution in the QRDR of GyrA and/or ParC was considered to confer resistance to fluoroquinolones. Besides, a total of 157 PMQR genes and chromosomal efflux pump (OqxA-B) were identified, 11 being unique (Supplementary Figure 8). OqxA-B was not considered to confer resistance to fluoroquinolones, while Qnr was (except for chromosomal Qnr in *C. freundii* (33) and *S. marcescens* (34)). A correct inference of susceptibility or resistance was obtained in 99.5% (186/187) strains (Figure 2). The only ME observed was in strain *E. cloacae* 7 that had a serine to threonine substitution at GyrA83. Although this position is a hotspot for mutations leading to fluoroquinolone resistance, the conservative nature of the mutation is not expected to modify the phenotype.

## Discussion

The main result of this study is the overall high rate of correct predictions of antibiotic susceptibility from WGS data when predictions were attempted, with a 97.6% concordance rate between phenotypic testing and genotypic results. The results obtained in non-AmpC strains were similar to those already published for *E. coli* (8–11) and *K. pneumoniae* (9, 13), and so were the results obtained with AmpC-producing strains when the inference was attempted. However, the prediction of the overexpression of AmpC from genomic data will require more data in order to be effective. Indeed, we found that only a minority of AmpC-producing strains had *ampD* and *ampR* genes sharing a high identity with that of the reference genome, which highlights the need for sequencing more strains from AmpC-producing species in order to cover their genetic diversity.

For amikacin, WGS could even be more accurate in detecting resistance than conventional testing. The EUCAST guidelines recommend to interpret as non-susceptible to amikacin a strain which is resistant to tobramycin but apparently susceptible to gentamicin and amikacin, i.e. suggesting the production of an AAC(6’)-I enzyme (35). Nonetheless, the application of this recommendation is compromised by the co-production of an AAC(3)-II) which confers resistance to gentamicin (13 strains in our set) and AAC(6’)-I – producing strains may falsely appear to be susceptible to amikacin. Indeed in our study, we observed 13 MEs for amikacin-susceptible phenotype associated with the presence of an AAC(6’)-I. In this case, the detection of ARGs and the subsequent phenotype interpretation could be more efficient than conventional testing. Besides, we found no errors in the prediction of susceptibility to gentamicin.

We observed a total 23 VMEs in our set of 187 strains. VMEs may cause a failure in the treatment of patients since antibiotics inferred to be active on a pathogenic strain may have no effect. The majority of VMEs occurred for beta-lactams and could be tentatively attributed to an overexpression of chromosomal genes (*bla*_SHV_ in *K. pneumoniae, bla*_OXY_ in *K. oxytoca, bla*_HugA_ in *Proteus vulgaris, bla*_AmpC_ and loss of permeability in *E. cloacae*), which addresses the limits of WGS when addressing the levels of expression of ARGs. However, when the increased gene expression is due to gene multiplication, the use of long reads can possibly resolve this issue (36). Otherwise, we need more data linking mutations to the expression of ARGs and mutations affecting the expression of intrinsic genes such as *bla*_AmpC_, or to use naive methods such as machine learning-based methods (15). To the best of our knowledge, the quantification of the expression of genes (e.g. via transcriptomics) for inferring antibiotic susceptibility has not been tested yet for Enterobacteriaceae. One VME was due to the incompleteness of the ResFinder database (absence of the class A beta-lactamase *bla*_HMS-1_ sequence). The VME in *P. mirabilis* 75 was left unexplained. Finally, the *E. coli* strain 53, which produced a CMY-42 and an OXA-181 was resistant to cefepime. CMY-42 is a derivative of CMY-2 with a higher activity on cefotaxime and ceftazidime, but its activity on cefepime remains unclear (37). Of note, the chromosomal *bla*_AmpC_ of the strain was wild type (data not shown). The decreased susceptibility to cefepime of *E. coli* 53 could then be explained by the combination of the CMY-42 and OXA-181, the latter having a moderate activity against cefepime (38)).

Our study has limitations. We analyzed a limited number of strains per species, and our panel may not be representative of the every-day situation in clinical laboratories. We focused on various resistance profiles for *E. coli* and *K. pneumoniae*, while more wild-type strains were included for other species such as *Citrobacter koseri* and *P. vulgaris*. The predictive values of WGS for inferring susceptibility indeed depend on the local epidemiology and they were not calculated here. Also, we only considered the antibiotics given in the early stages of HAP, and furthermore we identified situations where we chose not to infer susceptibility from genomic data. Hence, we acknowledge that for some species-antibiotic combination, the genotype-to-phenotype inference may not currently performant due to intrinsic limitations of WGS, but that in the majority of situations the inference could be attempted with good performances.

Clinical metagenomics is expected to reach diagnostic laboratories in the coming years. In the context of HAP, clinical metagenomics has the potential to decrease the turn-around time from sample to results and to definitive antibiotic therapy, potentially providing higher chances of cure or a better outcome. Hence, we chose to consider only a limited set of antibiotics – those used in the context of HAP treatment. We also chose to avoid inference attempts when available data would not support a strong prediction (essentially due to gene expression unpredictability), leaving some uncertainty for some species-antibiotic combination. From this clinical perspective, we observed very good performances for ceftazidime (in non-AmpC producing strains), cefepime (all strains), fluoroquinolones and aminoglycosides, with rare VMEs in our set. Our results support that clinical metagenomics could allow a reliable antibiotic susceptibility prediction for relevant antibiotics in the HAP context, provided that a good genome coverage of the bacterial pathogen would be achieved.

## Acknowledgements

This work was funded by bioMérieux. The authors are grateful to the bioMérieux/HUG HAP/VAP metagenomic group for helpful discussions: Gaspard Gervasi, Ghislaine Guigon, Maud Tournoud, Sébastien Hauser, Emmanuelle Santiago-Allexant, Véronique Lanet, Alex van Belkum, Nadia Gaïa and Damien Baud.

